# Parallels of quantum superposition in ecological models: from counterintuitive patterns to eco-evolutionary interpretations of cryptic species

**DOI:** 10.1101/2022.01.20.477053

**Authors:** David G. Angeler, Hannah B. Fried-Petersen

## Abstract

Superposition, i.e. the ability of a particle (electron, photon) to occur in different states or positions simultaneously, is a hallmark in the subatomic world of quantum mechanics but non-sensical from the perspective of macro-systems such as ecosystems and other complex systems of people and nature. Using time series and spatial analysis of bird, phytoplankton and benthic invertebrate communities, this paper shows that superposition can occur analogously in redundancy analysis (RDA), a form of canonical ordination frequently used by ecologists. Specifically, we used correlation analysis to show that species can be associated simultaneously with different orthogonal axes in RDA models, a pattern reminiscent of superposition. We discuss this counterintuitive result in relation to the statistical and mathematical features of RDA and the recognized limitations with current traditional species concepts based on vegetative morphology. We suggest that such “quantum weirdness” is reconcilable with classical ecosystems logic when the focus of research shifts from morphological species to cryptic species that consist of genetically and ecologically differentiated subpopulations. We support our argument with theoretical discussions of eco-evolutionary interpretations that should become testable once suitable data are available.

## Introduction

The laws of quantum physics describe reality at the level of atoms and subatomic particles (see Rae [2004] and Kastner [2015] for non-specialist treatments of quantum physics). These laws differ radically from those of classical physics that characterize systems at the macroscopic level, such as ecosystems and other complex systems of people and nature. Quantum physics defies the logic that underpins the reality accessible to our human senses, and both branches of physics are therefore generally perceived as incompatible and mutually exclusive. However, there is increasing evidence that the ways the brain (Wendt 2015), human societies (Zohar and Marshall 1995), and economic systems (Orrell 2018) work is often reminiscent of quantum physical phenomena. This highlights synergies that can be exploited for genuinely novel interdisciplinary research (Bull and Gordon 2015).

Ecologists have already begun to use physics as an analogous model for describing patterns and processes in ecosystems such as species diversity and distribution modeling (Rodríguez et al. 2015a; Real et al. 2016), the quantification of evolutionary processes (Rodríguez et al. 2015b), dynamic management and conservation of nature reserves (Bull 2015), modeling of desertification (Bagarello 2019), and management for sustainability (Alrøe and Noe 2016). Inspired by Erwin Schrödinger’s famous cat that is simultaneously dead and alive (Schrödinger 1935), work has also envisioned the potential to create quantum superposition (i.e. particles occur at several places or in different states at the same time) experimentally in viruses and microorganisms such as tardigrades (Romero-Isart et al. 2010). While the experimental induction of superposition states for living organisms is currently elusive, this paper will show and discuss the widespread but hitherto ignored occurrence of superposition in statistical modeling of plant and animal communities frequently used in ecology.

In this paper superposition is used in the form of an analogy, a useful approach for relating quantum mechanics with classical systems (e.g., Zohar and Marshal 1993; Orrell 2018). While the analogous use of superposition does not allow for a mechanistic 1:1 translation and application of superposition in the ecological models discussed here, it provides at least two opportunities for research. The first relates to a heuristic role for associative research that can spur surrogative learning and inductive thinking (Magnani et al. 1999, 2002; Swoyer 1991). Following from this potential, the second relates to the generation of new questions and hypotheses (Hesse 1974; Holyoak and Tagard 1995). This paper exploits both opportunities, using redundancy analysis as a heuristic of superposition in ecological models, followed by a novel theoretical discussion about eco-evolutionary implications, particularly pertaining to cryptic species.

Redundancy analysis (RDA) (Rao 1964; van den Wollenberg 1977) is a multivariate constrained ordination technique that is routinely applied by community ecologists. RDA has been often used to assess the distributions of freshwater, marine and terrestrial species assemblages as a function of environmental variables (Legendre and Legendre 2012; Borcard et al. 2018). RDA has also been used extensively in time series, spatial and combined spatial-environmental (variance partitioning) analyses. These analyses model temporal or spatial patterns in ecological data using mathematical eigenvector representations of time and space in the analysis (Baho et al. 2015; Peres-Neto et al. 2006; Dray et al. 2006, 2012). RDA assesses the variation in a set of response variables, such as species, that can be explained by a set of explanatory variables (e.g. environmental variables, spatial coordinates, or a time vector). More specifically, RDA synthesizes linear associations between components of response variables that are “redundant” with, or “explained” by a set of explanatory variables (Buttigieg and Ramette 2014). This occurs in the form of creating orthogonal (statistically independent) canonical axes (or RDA axes) that are built from linear combinations of response variables that are simultaneously linear combinations of the explanatory variables.

The RDA axes resolved by the models have often been correlated with the time series of individual taxa (Baho et al. 2014). The result can be a classical “either/or scenario” in which a species correlates significantly with one or the other axis (Figure 1A). Due to the nature of RDA, a situation may arise where a taxon will be significantly correlated with more than one axis. Using the superposition analogy from quantum physics a paradox arises: the species occurs in different places at the same time, or in the context of the models, simultaneously at different RDA axes, manifested in a “both/and scenario” (Figure 1A). Such a result is counterintuitive from the perspective of the logic of classical ecology, and macro-systems in general. Such an “absurdity” may explain the neglect of this feature of RDA by ecologists.

**Figure 1:**
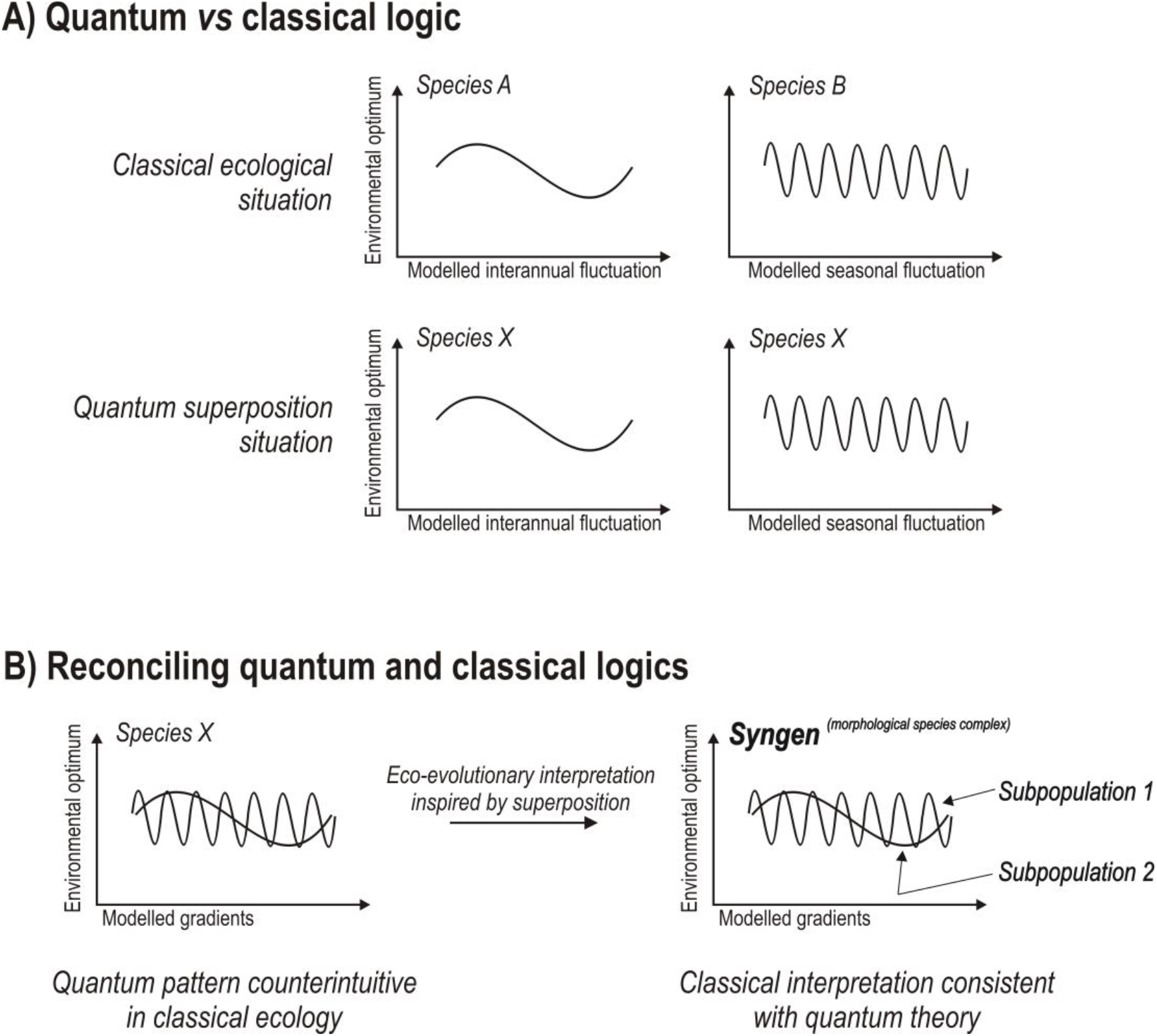
Schematic representing A) a classical and “quantum superposition” situation in RDA models. The classical situation represents an “either/or scenario” that arises when a species is associated with a specific gradient or RDA axis (e.g. species A with nutrients, species B with temperature). The quantum superposition scenario comprises a “both/and scenario” which emerges when a species is simultaneously associated with both gradients, as is the case with species X (see text). B) represents a reconciliation of quantum and classical logics in an attempt to inspire and advance novel eco-evolutionary theory of cryptic species or syngens.

The aims of this paper are two-fold. First, we capitalize on such “quantum weirdness” in RDA models building on the rationale that classical analyses are not necessarily incompatible with demonstrating quantum physical features (Orrell 2018). For this purpose, our analogous use of quantum features builds on an adaptation of our ecological analyses to better fulfill the premises of quantum theory. That is, we eliminate linear trends in time series (Baho et al. 2015) to create a quantum analogue of the classical RDA models. This approximation can be achieved through detrending the RDA models. This simulates quantum systems that deal with time less stringently than classical systems, meaning that the future can inform the past, which has been demonstrated in delayed-choice quantum erasure experiments (e.g. Kim et al. 2000). Concomitantly, the RDA approach can emulate the particle aspect (e.g. electrons and photons) of the particle-wave duality in quantum systems, through analysis of species presence/absence patterns rather than biomass- or abundance-based data.

The second goal of this paper goes beyond the mere demonstration of analogous patterns of superposition in RDA models. It is increasingly recognized that the adoption of quantum analogies in classical systems has potential to inspire novel interpretations of their dynamics that are not at odds with classical logic (Wendt 2015, Orrell 2018). In the broadest sense, this suggests that the logics of a quantum and classical world are reconcilable and that ultimately current theories can be advanced through the cross-fertilization of both logics (Bull and Gordon 2015). This paper attempts such unification by showing that quantum superposition can inspire discussions about eco-evolutionary theories. Specifically, we will discuss ecological differentiation of reproductively isolated lineages of taxa that conserve morphological similarity (e.g. cryptic species or syngens; Sonneborn (1957)).

We envision that quantum superposition in RDA models is not at odds with classical interpretations when genetically and ecologically differentiated subpopulations rather than the morphological species per se are analyzed (Figure 1B). That is, differentiated subpopulations are expected to fit the “either/or” scenario of classical ecology in the models, relative to analyses based on morphological species that become “smeared out” across axes resulting in the paradoxical “both/and” scenario arising from superposition. We will discuss the ecological implications of these scenarios in relation to statistical features of RDA and recognized shortcomings of currently applied species concepts based on vegetative morphology (Fišer et al. 2018). Although the potential for reconciling classical and quantum logics for better understanding ecological systems may be enormous, our discussion about eco-evolutionary implications is purposefully preliminary, speculative and theoretical until suitable data for empirical testing become available.

## Material and methods

We selected data of different taxonomic groups and ecosystems and used time series and spatial modeling based on redundancy analysis to showcase superposition as a general allegorical feature in our modeling approach.

### Data

For time series analyses we used two data sets. The first data set was obtained from the publicly available US Breeding Bird Survey (BBS) of North America, which contains avian community composition that is collected by qualified observers along georeferenced, permanent roadside routes across North America (Sauer et al. 2017). Along each approximately 39.5 km route, observers make 50 stops once every 0.8 km and conduct point-count surveys. During each survey, observers record for three minutes the abundance of all bird species that are acoustically or visually detected within a 0.4 km radius. Surveys start thirty minutes before local sunrise and last until the entire route is finished. To increase uniformity in probability of bird detection, surveys are conducted only on days with little or no rain, high visibility, and low wind.

For this study, we selected the South Central Plains as an example of a terrestrial ecosystem. We averaged three transects spanning the latitudes 31.8 to 33.4, which were consistently sampled between 1968 and 2014 (47 years of data). We removed all aquatic species from the families Anseriformes, Gaviiformes, Gruiformes, Pelecaniformes, Phaethontiformes, Phoenicopteriformes, Podicipediformes, Procellariiformes, and Suliformes from analyses because of known negative observation biases for waterfowl compared with terrestrial avian families (Roberts et al. 2019ab). We also removed hybrids and unknowns, and we condensed subspecies to their respective species following Angeler et al. (2020).

Our second data set for time series analysis contains phytoplankton community data from lake Stensjön, a small (surface area 0.57 km^2^), nutrient-poor, circumneutral lake located in the northern boreal forest biome of central Sweden (long: 14.77, lat: 56.45). Lake Stensjön is included in the Swedish National Lake Monitoring Program, which was established in the 1970s to assess the impact and recovery of anthropogenic acidification (Fölster et al. 2014). The monitoring program is overseen and regulated by the Swedish Agency for Marine and Water Management (HaV: https://www.havochvatten.se/en). Data are open access and available: http://miljodata.slu.se/mvm/.

For this study we used data spanning the period 1992 to 2018. Integrated samples of phytoplankton were collected from 5 sites in the upper stratification layer of the lake (epilimnion) in August with a plexiglass tube sampler (2 m long, inner diameter 10 cm), pooled and preserved in Lugol’s solution. Phytoplankton counts were made using an inverted light microscope following the modified Utermöhl technique commonly used in Skandinavia (Olrik et al. 1989). Taxa were usually identified to the species-level taxonomic unit.

For the spatial analysis we used an exhaustive set of littoral invertebrate community data from 105 lakes sampled in 2017 that were distributed across Sweden (Fried-Petersen et al. 2020; Figure 1). The studied lakes all belong to the Swedish National Lake Monitoring program (see above), are medium sized (area = 0.03–14 km^2^, mean = 1.5 km^2^) and are considered least disturbed in terms of no impact from point sources of pollution and land-use (Fölster et al. 2014). Sampling and analyses protocols for invertebrates are certified and quality controlled through the Swedish Board for Accreditation and Conformity Assessment (SWEDAC; http://www.swedac.se/en/) and followed Swedish standards (SS‐EN 27828).

Invertebrates were collected from each lake in one wind‐exposed, vegetation‐free littoral habitat during late autumn. In the most northern lakes, sampling was conducted between September and November to achieve similar seasonal conditions across surveys. Five replicate samples were taken, using standardized kick sampling with a hand net (0.5 mm mesh size). For each sample, the bottom substratum was disturbed for 20 s along a 1 m stretch of the littoral zone at a depth of ~0.5 m. Invertebrate samples were preserved in 70% ethanol (estimated final concentration) in the field and processed in the laboratory by sorting against a white background with 10× magnification. Invertebrates were identified to the finest taxonomic unit possible and counted using dissecting and light microscopes.

### Modeling and analyses

#### RDA analyses

We carried out two time series analyses, one for the bird and the other for the phytoplankton data set, and one spatial analysis for the invertebrate data set based on RDA. All statistical analyses were carried out in R 3.6.1 (R Development Core Team, 2019) using packages vegan (Oksanen et al. 2019), adespatial (Dray et al. 2019), ade4 (Dray and Dufour 2007) and quickMEM (Borcard 2016).

For the time series analyses, Moran Eigenvector Maps (MEM) (Dray 2019, Dray et al. 2006, 2012), which comprise a set of orthogonal temporal variables, were obtained through the conversion, akin to a Fourier transformation, of the time vectors of the bird and phytoplankton time series. These time vectors consisted of 47 steps (sampling years) between years 1968 and 2014 for birds and 24 steps between 1992 and 2018 for phytoplankton, respectively. As a result of the Fourier transformation, these temporal MEM variables take on the shape of sine waves of different wavelengths, which allows assessing fluctuation patterns at different inter-annual and interannual scales in the bird and phytoplankton data. These MEM variables are then used as explanatory variables to model temporal relationships in the bird and phytoplankton incidence data using redundancy analysis (RDA) (Baho et al. 2015).

Using forward selection, RDA selects significant MEM variables that best explain the temporal structures extracted from the bird and phytoplankton species matrices. The modeled temporal patterns that are extracted from the data are collapsed onto significant RDA axes, which are tested through permutation tests. The R software generates linear combination (lc) score plots, which visually present the modeled temporal patterns that are associated with each RDA axis. That is, individual RDA axes indicate fluctuation patters at temporal frequencies or scales that are statistically, and presumably ecologically, independent from those of other axes (i.e. orthogonal canonical axes). In the context of this study, we consider the different axes resolved by RDA as analogues of different quantum states existing simultaneously. More concretely, this analysis allows us to identify bird and phytoplankton taxa showing different temporal patterns at the same time (i.e. a form of “temporal superposition”; see below). All bird species raw-abundances averaged from three transects and phytoplankton biovolume data were transformed into presence-absences prior to the analysis and models were detrended when monotonic patterns of change were identified.

The spatial analysis using invertebrates implemented the same analysis steps as the time series analysis, with the difference that spatial coordinates comprising longitude and latitude of the sampling locations in the 105 lakes were used for constructing spatial MEM variables. As a result, the graphical representation of the spatial analysis presents significant RDA axes in the form of two-dimensional spatial planes instead of one-dimensional temporal plots, as is the case with the time series (Figure 2). These independent spatial planes were considered analogous of co-existing independent spatial domains at which invertebrate species might occur simultaneously, indicating spatial superposition.

**Figure 2:**
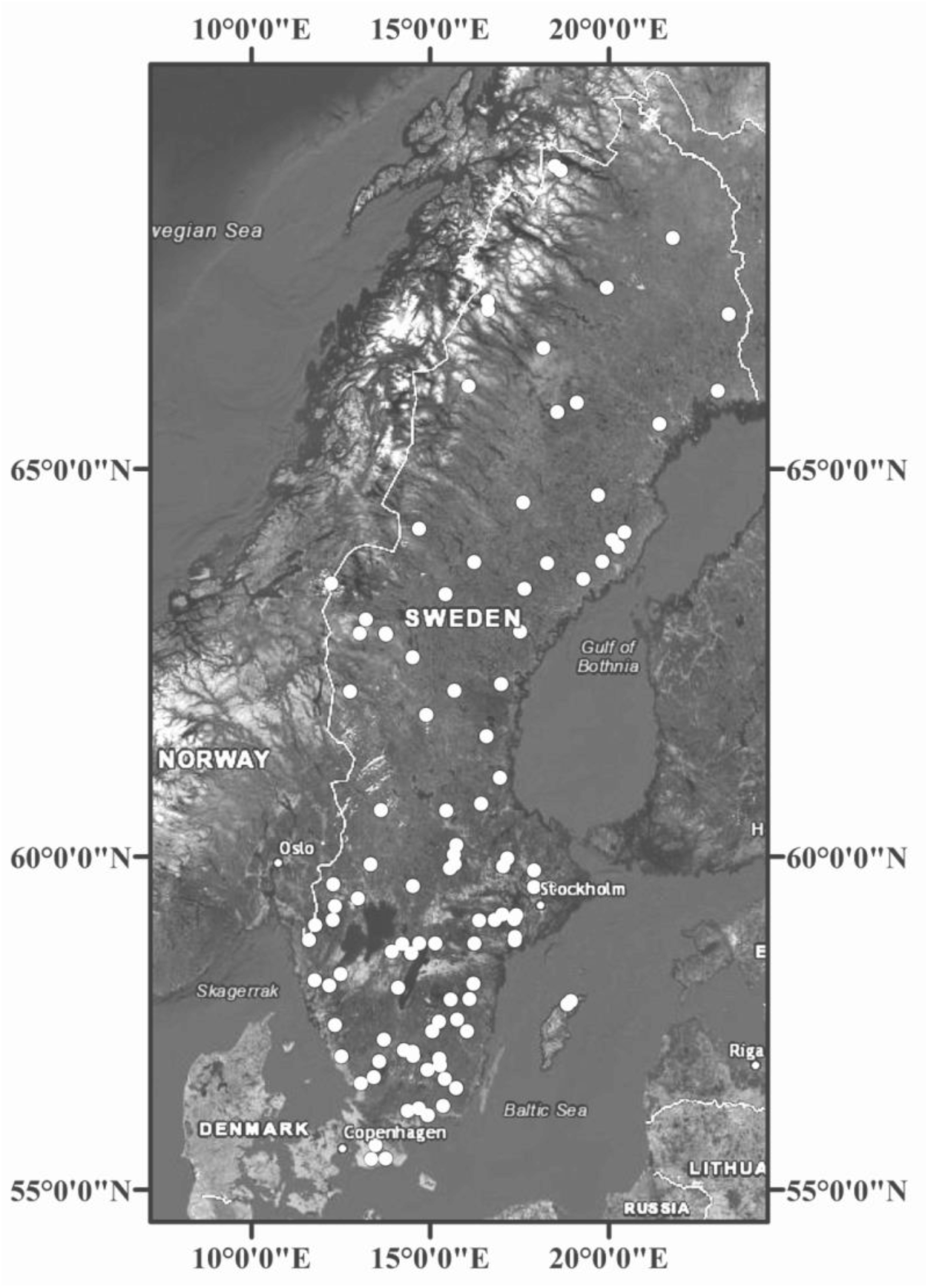
Map of Sweden showing the distribution of 105 lakes (white dots) used for the spatial analysis of littoral invertebrate data.

#### Correlation analyses

The RDA analyses formed the foundation to test for superposition through the identification of statistically independent objects (distinct temporal and spatial patterns associated with orthogonal canonical RDA axes). In this analysis step, we formally assessed whether bird, phytoplankton or invertebrate data significantly correlate with one or more of the identified axes. To infer quantum superposition of species in the models significant correlations with more than one axis are prerequisite. Following Angeler et al. (2015), we used Spearman rank correlation analysis to relate incidence data of individual bird, phytoplankton and invertebrate taxa with the modeled patterns (lc scores) associated with the RDA axes of the respective models. This allowed us to assess how prevalent superposition is in the analyzed communities relative to correlations of species with only a single or no axis across species and data sets. Taxa that do not show significant correlations are considered to be stochastic because their dynamics are unrelated to the deterministic gradients revealed by RDA and are thus random with respect to these specific analyses (Baho et al. 2014; Angeler et al. 2020). However, such species have traditionally been down-weighted by ecologists using RDA, although they may be relevant for understanding important ecological facets such as adaptive capacity or resilience (Angeler et al. 2019). We report the prevalence of all these fractions (species correlating with more than one axis [i.e. those showing superposition], those correlating with only one axis, and those not correlating with any axis) for broadest contextualization and comparisons of our results.

## Results

### RDA models

Time series analysis for birds and phytoplankton and spatial analysis for invertebrates revealed significant models for all organism groups, although the proportion of variance of the minimum models (adjusted R^2^) explained was low (birds: 0.08, phytoplankton: 0.2, invertebrates: 0.04). All models resolved more than one significant temporal or spatial dimension (RDA axes), thereby building the necessary basis for testing for the “both/and” superposition scenario. These models were manifested in 2 and 4 significant temporal dimensions for birds and phytoplankton, respectively; the spatial model for invertebrates revealed two significant spatial patterns (Figure 3). The time series models for birds and phytoplankton showed broader scale (i.e. slower) patterns of community fluctuations associated with RDA 1 relative to the other RDA axes, which displayed faster community turnover (Figure 3). Similar patterns were found in the spatial analysis for invertebrates, which displayed slightly more broad-scale patterns with RDA 1 relative to RDA 2.

**Figure 3:**
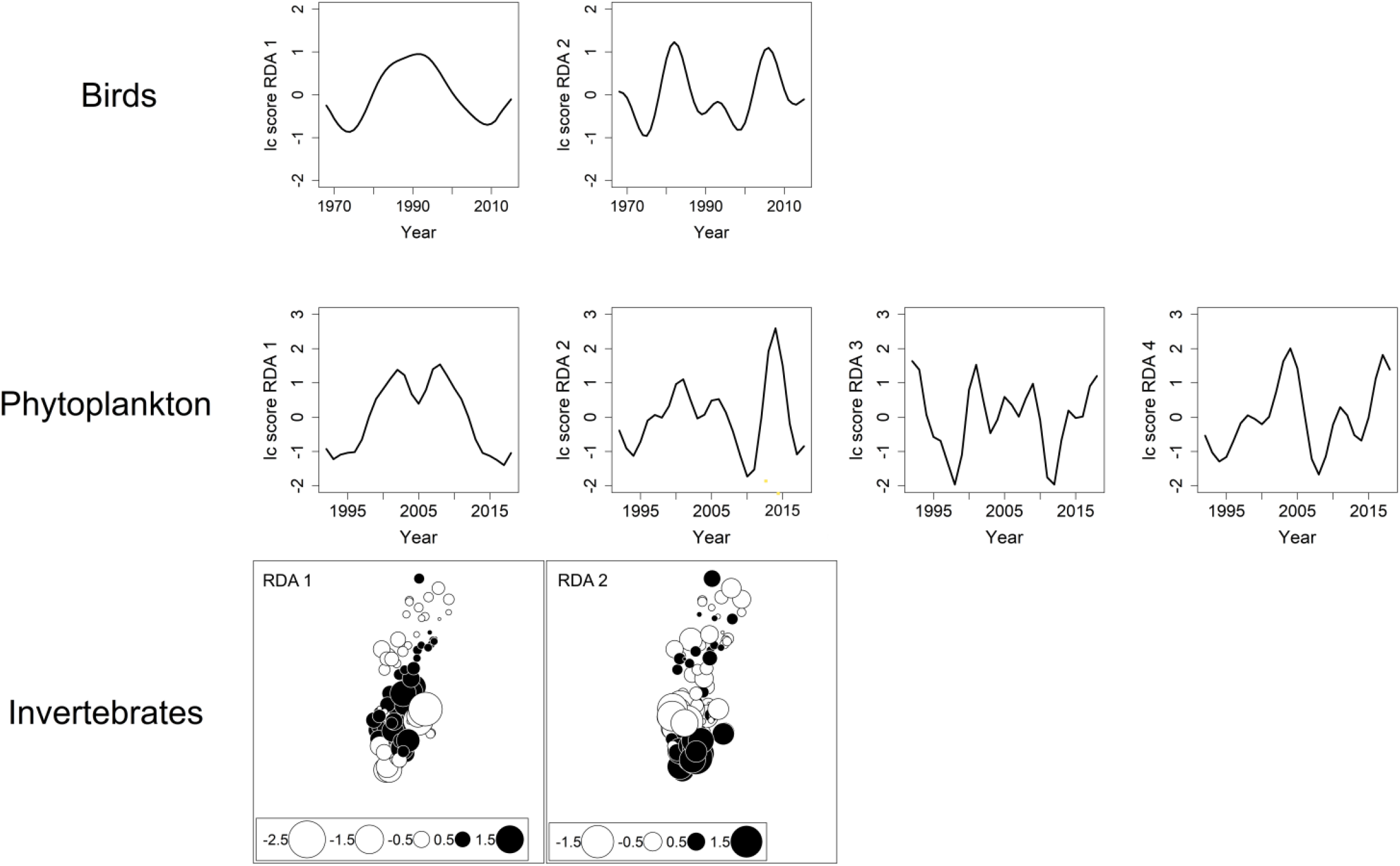
Linear combination score plots associated with significant RDA axes in the MEM-RDA analyses showing statistically independent temporal patterns for birds and phytoplankton and spatial patterns for littoral invertebrates.

### Correlation analyses

Spearman rank correlation analyses revealed a low prevalence of superposition in the RDA models. For birds, 5 species (4.5% of the total number of species (n = 110)) correlated with both significant axes of the RDA model, thus showing superposition in terms of simultaneously displaying independent temporal dynamics (Table 1). The remaining species correlated either with RDA 1 (9%) or RDA 2 (4.5%) or were not significantly correlated with any axis (stochastic species) (82%). For phytoplankton, 21 species (7% of the total number of species (n = 310)) showed superposition (Table 1). The remaining species correlated either with RDA 1 (11%), RDA 2 (6%), RDA 3 (4%) or RDA 4 (3%), or were not significantly correlated with any axis (stochastic species: 68%). For invertebrates, 3 species (3% of the total number of species (n = 119) showed superposition in terms of occurring simultaneously in the two spatial dimensions (planes) resolved by the RDA (Table 1). The remaining species correlated either with RDA 1 (26%) or RDA 2 (6%) or were stochastic species (66%).

**Table 1:**
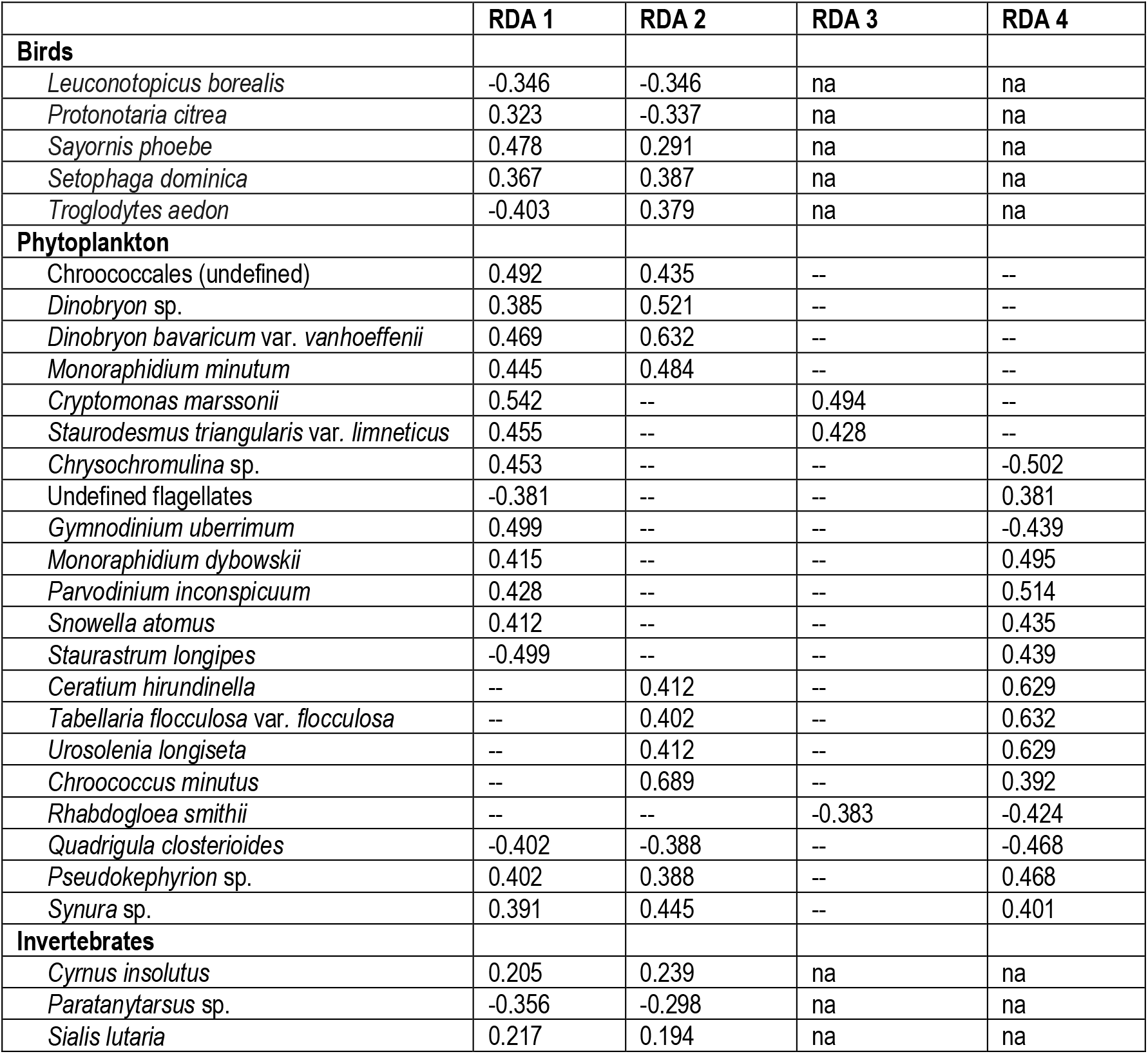
Results from Spearman rank correlation analysis testing for superposition. Shown are significant spearman rank correlation coefficients (rho) at a significance level P ≤ 0.05. Abbreviations: na, not applicable; -- , not significant. Note that signs of correlations are not relevant for assessing superposition.

## Discussion

The results of this study support our first goal showing that quantum superposition occurs in analogous form in RDA models across distinct organism groups and ecosystems. However, the prevalence of superposition was low. This may be due to intrinsic limitations in the approach and its adaptation to test for quantum superposition. An analytical approach as the one used in this paper frequently yields limited explanatory power due to the nature of correlative analysis in which residual variation can be introduced due to the accumulation of noise resulting from sampling, survey designs, ecosystem history and system-intrinsic variation (Leibold et al. 2010). The low amount of variance explained can also be attributed to the correction of R^2^ - values by the number of explanatory variables for obtaining appropriate models (Peres-Neto et al. 2006). Furthermore, detrending models is an additional source of loss of variance explained (Borcard et al. 2004; Baho et al. 2015). Finally, using incidence rather than quantitative data of species (abundances, biovolumes) in the RDA models and correlation analyses might have further contributed to a decreased statistical performance of our analysis and thus the detection of low prevalence of superposition. As a result the estimates of prevalence of superposition in our study might be conservative. However, the overall partitions of species correlating with a single or no axis (stochastic species) matches results from previous times series (Baho et al. 2014) and spatial analysis (Angeler et al. 2015), which suggests that incidences of superposition in RDA models might be generally low. Notwithstanding, we acknowledge that a different analysis design could have probably resulted in different prevalence patterns of superposition but we purposefully traded off statistical performance in favor of adapting the approach to specifically account for premises of quantum mechanics (“free flow of time”, species as particles) in our analyses. Also, in the present context, and from the biological side, we can currently not infer why the specific taxa in our analyses showed superposition, or the factors that explain these patterns. This warrants future research.

Our study is based on the analogous use of quantum superposition in the RDA models. Using such an analogy clearly prevents a mechanistic 1:1 extrapolation, application and interpretation of superposition in ecology. In quantum mechanics, superposition arises from the simultaneous occurrence of different states of an object. These states comprise potentialities in an intangible reality (Kastner 2015) and only one of these potentialities will eventually actualize and manifest upon measurement while all other potentialities become “annihilated”, according to the Copenhagen interpretation of quantum physics. In a simple example, an electron can simultaneously have a left and right spin before measurement, but once measured either the left or right spin will become apparent. Superposition, while being the norm for subatomic entities is therefore clearly at odds with how we generally understand and interpret macroscopic situations. That is, individuals in ecological communities are manifested, tangible and measured entities that have been subjected to sampling, quantification and data analyses rather than inexistent, unperceived potentialities. The analogy in this study therefore, rather than comprising superposition in the sense of quantum physics, builds on a pattern akin to superposition that is generated by the RDA models. In this specific approach, this is due to the linear associations between the species that are redundant with and explained by a set of temporal or spatial predictor variables. The patterns of superposition of species as measured entities in the RDA models therefore results entirely from a mathematical and statistical procedure. That is, from creating orthogonal RDA axes that are built from linear combinations of species and explanatory variables (Buttigieg and Ramette 2014).

Despite the discrepancies between the manifestation of superposition in quantum mechanics and in the RDA models, the analogous use of superposition for the purpose of this study is useful for stimulating research at the intersection between quantum mechanics and classical ecology, as has been shown for social (Zohar and Marshall 1993) and economic (Orrell 2018) systems. This brings us to the second goal of this study: reconciling the manifested quantum phenomena in the RDA models with eco-evolutionary patterns that are consistent with classical ecological logic. We will base our discussion on genetically differentiated subpopulations of taxa with near-identical morphology, i.e. cryptic species or syngens (Sonneborn 1957).

Unique vegetative morphology can independently emerge at different times during evolution, showing that morphology is not necessarily a marker of a monophyletic group or taxonomic species (Angeler et al. 1999). That is, species which share vegetative morphology (i.e. “morphological species”) can consist of reproductively isolated and ecologically differentiated subpopulations. Only members of a specific subpopulation (cryptic species or syngens) are compatible for mating, thereby fitting the biological species concept (Mayr 1948; Manhart and McCourt 1992). Cryptic species are remarkably diverse among microscopic organisms (e.g. Fenchel et al. 1997; Schagerl et al. 1999), but are also widespread in animals and plants (e.g. Baker and Bradley 2006; Fernandez et al. 2006; Bickford et al. 2007; Pfenninger and Schwenk 2007; Shneyer and Kotseruba 2015). With the continued development of molecular techniques even more cryptic species across organism groups, ecosystems and biomes are likely to be discovered. Despite the potential arising for biodiversity research, ecological and evolutionary factors that shape or are shaped by cryptic species have received limited research attention (Fišer et al. 2018). Given this dearth of information in the literature and the lack of data for empirical testing the following discussion about the reconciliation of our results with quantum theory is theoretical and aimed at stimulating future research.

Fišer et al. (2018) consider cryptic species as a window for a paradigm shift of the species concept. Our results suggest that such a consideration is warranted. Ecological research is strongly biased towards morphological species routinely evaluated in traditional taxonomic studies, rather than using cryptic species complexes concealed in a morphological species identified by molecular methods and experiments that ascertain their ecological distinctness. RDA has strong potential to identify groups of taxa with similar ecological patterns resulting from intrinsic (e.g. nutrients, temperature, biological interaction) and extrinsic (e.g. habitat connectivity, spatial patterns) factors and temporal change that affect ecological communities. The multiscalar nature of RDA also allows distinguishing between different deterministic patterns and can therefore indicate ecologically distinct groups of species in a community. RDA therefore has strong potential to test explicitly for the complex and non-linear factors that shape ecosystems, including cryptic species complexes, across different scales of space and time (Holling 1992; Allen et al. 2006, 2014; Nash et al. 2014)

While a wealth of different study and analysis designs may have the potential to reveal the ecological distinctness of cryptic species, the usefulness of RDA per se as a quantitative method for assessing such ecological differentiation of cryptic taxa needs further evaluation. However, our study suggests that RDA serves as a useful heuristic, inspired by quantum physics, to inform about the limitations when not accounting for ecological differentiation of subpopulations or when relevant environmental factors are not included in the analysis. Specifically, using traditional taxonomic surveys based on vegetative morphology may mask the differentiation of cryptic species in the analysis and add noise. The heuristic value of RDA therefor resides in indicating shortcomings of traditional analyses in the form of morphological taxa becoming “smeared out” across RDA axes, resulting in the counterintuitive and illogical superposition pattern. This quantum superposition becomes allegorical of the limitations of species concepts based on morphological criteria reported in the literature (Fišer et al. 2018).

There is increasing evidence that cryptic species not only differ at the genomic level but also in environmental optima mediated by different functional traits (Bailet 2021). This suggests ecological distinctness and the occupation of different ecological niches, which may manifest with the association of cryptic species with different axes in the RDA heuristic (Figure 1b, right panel). We currently lack the exhaustive genomic functional trait data with exhaustive spatial and temporal resolution for testing to what extent the eco-evolutionary analysis of cryptic species fits our heuristic. Future research using such extensive data sets may be useful for this purpose. The allegorical us of quantum physics in such research may inspire hitherto unrealized potential for novel research.

## Acknowledgments

We thank Richard K. Johnson for critical reading of the manuscript.

## Author contributions

DGA conceived the study and wrote the paper. HBF-P analyzed the data, prepared the figures and commented on an advanced paper draft. Both authors have approved submission of the paper.

## Data and script availability

Published data and R code can be found in the Zenodo archive at https://doi.org/10.5281/zenodo.5880132.

